# The costs of global protected-area expansion (Target 3 of the post-2020 Global Biodiversity Framework) may fall more heavily on lower-income countries

**DOI:** 10.1101/2022.03.23.485429

**Authors:** A. Waldron, C. Besancon, J.E.M. Watson, V.M. Adams, U.R. Sumaila, S.T. Garnett, A. Balmford, S.H.M. Butchart

**Affiliations:** Cambridge Conservation Initiative, David Attenborough Building, Pembroke St, Cambridge CB2 3QZ, UK; Working Ant Consultancy Cambridge Ltd., Cart House 2, Copley Hill Business Park, Cambridge Rd, Cambridge CB22; Global Park Solutions, 2809 Duncan Drive, Missoula Montana 59802 USA; School of Earth and Environmental Sciences, University of Queensland, St Lucia 4072 Australia; School of Geography, Planning, and Spatial Sciences, University of Tasmania, Hobart TAS 7001 Australia; Institute for the Oceans and Fisheries and the School of Public Policy and Global Affairs, University of British Columbia, Canada; Research Institute for Conservation and Sustainable Livelihoods, Charles Darwin University NT 0909 Australia; BirdLife International, David Attenborough Building, Pembroke St, Cambridge, CB2 3QZ, UK

## Abstract

One of the biggest stumbling blocks for global environmental agreements is how higher-income and lower-income countries share the costs of implementing them. This problem has become particularly acute as biodiversity and climate ambitions have increased across recent COPs (Conferences of Parties). Here, we estimate the likely distribution of costs for one of the most ambitious proposals: draft Target 3 of the Global Biodiversity Framework (GBF), which would increase coverage of protected and conserved areas (PCAs) to 30% of global land and sea area - more than triple the current value. Since the GBF does not specify where new PCAs would be placed, we use three scenarios of how Target 3 might be implemented, cost those scenarios, and then compare the mean distribution of costs across World Bank income groups. We find that in relative terms, lower-income countries could face considerably larger financial burdens than high-income countries, even though the benefits of conservation are disproportionately enjoyed by high-income countries. Lower-income countries would also face larger increases in the amount of land or sea under conservation, implying higher opportunity and establishment costs. Resolving this potential cost-sharing inequity may be a key requirement to achieve consensus on draft Target 3, and indeed on ambitious environmental proposals more generally.

---

One of the key proposals in the current draft of the Convention on Biological Diversity’s post-2020 Global Biodiversity Framework is to conserve 30% of both land and sea for biodiversity by 2030 (draft Target 3)^1,2^. Achieving this target would help mitigate both the biodiversity crisis and the land-use element of the climate crisis, bringing long-term economic benefits locally and globally^3^. However, meeting the target will also require substantial increases in the area currently under conservation and therefore, in financial costs. The experience of previous negotiations suggests that the way these costs become distributed across countries (and whether that distribution is perceived as equitable) risks being a major sticking point. For example, lower-income countries (sometimes referred to as developing countries or the Global South) may perceive that they are being asked to make financial commitments they view as unfair or beyond their economic capacity. Conversely, high-income countries may object to requests for international financial assistance (e.g. biodiversity aid^4^) that they regard as excessive or poorly justified. A fair cost distribution is not only a moral issue, but also a pragmatic and political one: high-income countries may need to offer sufficiently attractive conditions to lower-income countries to secure the latter’s cooperation on global environmental issues^5^. This consideration is particularly relevant to the CBD and UNFCCC because lower-income countries harbour much of the earth’s remaining biodiversity and carbon-rich natural habitat^6^, making their political assent critical.

The unprecedented ambition of the 30% target reflects the growing severity of the environmental crises. However, a large increase in ambition also magnifies the cost and with it, the level of political tension around any perceived distributional inequity. If the cost-sharing issue is not satisfactorily resolved^7^, there is a danger that the language of the final agreement will become diluted and vague, or that the political commitments made will lack the funding needed to adequately implement them. Underfunded “paper commitments” are unlikely to resolve the global environmental problems they claim to address, just as “paper parks^8^” fail to protect nature on the ground.

This danger is particularly acute when there is no information on how a target’s costs are likely to be distributed. Without such information, negotiations can be bogged down by what each party imagines (or fears) will happen, which makes consensus difficult. Here, therefore, we generate estimates of the likely geopolitical distribution of Target 3 costs (specifically, the likely increase in recurrent annual costs). We also analyse the target’s “territorial costs”: the amount of additional land and sea that countries might be asked to commit to conservation. Territorial costs are important because a larger national conservation area can imply larger opportunity costs3,9 and creation costs, in addition to the increase in recurrent annual costs. Noting that draft Target 3 calls for conservation of 30% of land and sea area through both protected areas and other effective area-based conservation measures (OECMs), recognising the importance of rights-based governance (not least because of the importance of Indigenous Peoples and Local Communities in achieving post-2020 goals^10^), we use the portmanteau term “protected and conserved areas” (PCAs) rather than the term “protected areas”.

A common point of political interest is how costs will be shared across economically wealthier and poorer countries. We therefore disaggregate our estimates of the Target 3 costs using the four World Bank income categories: High Income Countries (HICs), Upper-Middle Income Countries, Lower-Middle Income Countries, and Low Income Countries^11^. We first compare the HICs to all non-HIC countries (hereafter called LAMICs i.e. “low and middle income countries”); make a second comparison between HICs and LMICs (the group comprising Low Income Countries and Lower-Middle Income Countries only); and then report results for each individual World Bank Income category separately.

## Estimating geopolitical cost distributions

Although draft Target 3 specifies that expanded PCA systems should cover “especially areas of particular importance for biodiversity”^2^, the precise locations to be protected or recognised as OECMs in each country will be determined by individual governments. Estimates of costs and cost distributions are particularly useful during negotiations before the target itself is adopted or the new PCA locations known. Paradoxically, however, the costs depend on where the new PCAs will be located. To resolve this difficulty, we use three hypothetical, non-prescriptive 30% scenarios (taken from Waldron et al.^3^) as the basis for costing (see Methods). The scenarios follow the rubric of Target 3 but vary in reflecting a range of trade-offs between biodiversity importance and economic and political constraints. They are therefore designed to give a reasonable characterisation of the probable cost distribution, without being intended as defined blueprints for different solutions, nor as precise measures of the final cost once the details of any implementation have been decided. Scenario 1 identifies a set of new conservation areas (PCAs) with the pure goal of minimising global species loss; Scenario 3 takes a similar approach but adds the constraint that new PCAs cannot be placed upon land needed for future agricultural expansion; and Scenario 2 is intermediate between Scenario 1 and Scenario 3 (see Methods).

The main purpose of our study is to examine how the sharing of costs for Target 3 is broadly distributed across income groups. To explore the likely cost distributions, we calculate the mean recurrent annual costs across these three scenarios and disaggregate the results by income groups. Our main focus is the factor increase implied (i.e. how many times the current budget, or current amount of conserved national territory, has to be multiplied in order to achieve Target 3 under the scenarios), although we also show the modelled raw increases for information only. We additionally show the financial increase implied as %GDP, although we caution that this popular metric is not a good proxy of distributional equity across different national income levels (see below).

In many countries, PCA costs are already distributed across both domestic governments and international donors. When calculating “who-pays” cost distributions, one therefore needs to know what proportion of financing comes from domestic budgets and what proportion from international assistance (aid). We analysed the proportion of protected-area system budgets that is currently funded by aid across the different income groups in our data, then subtracted this from the current spending of LAMICs and added it to the current spending of HICs. We also used the aid data to estimate the likely impact of a doubling of aid on the distribution of costs for Target 3, by subtracting the (doubled) aid from the finance that LAMICs would need to provide, and then adding it instead to the finance that HICs would need to provide. In our results, we report the aid-adjusted distribution of costs under two conditions: (a) international aid remains at its current level, (b) international aid doubles. We note that our model can be used flexibly to estimate how any hypothetical increase in international financial assistance would affect cost distributions and their equitability.

## Likely distribution of cost burdens for Target 3

If aid remains at its current levels, lower-income countries would need to increase their PCA expenditure by a considerably larger multiple than HICs (see factor increases in Figure 1, Table 1). For example, on average (median value), HICs would need to increase their current expenditure by a factor of 1.77, LMICs by a factor of 5.41 and LAMICs by 5.53 (Table 1). Budget increases expressed as %GDP were typically low, at approximately 0.1% of GDP (Figure 1, Table 1). As the exception, the budget increase needed for low-income countries, the poorest income group, was disproportionately high at 0.32% of GDP, which is over three times the value for HICs, the wealthiest group (Figure 1). In raw dollar terms, HICs had the largest implied increase in expenditure (adjusted for aid): an increase of $12.5 billion, compared to $9.9 billion for LAMICs and $3.8 billion for LMICs (Figure 1, Table 1).

**Table 1.** The increase in annual financial costs and territorial costs implied by Target 3, per income group (mean across three potential scenarios after adjusting for aid). First value in each column shows result at baseline level of international assistance; second value models a doubling of international assistance to protected and conserved areas. Median increases take the median value for individual country outcomes; total increase takes the summed value for the income group as a whole and is therefore not directly comparable to the country medians. HIC = High-Income Countries, LMIC = Low-to-Middle-Income Countries (Low and Lower-Middle only), LAMIC = LMIC + Upper-Middle-Income Countries (i.e. all non-HIC countries), UpperMid = Upper-Middle Income Countries, LowMid = Lower-Middle Income Countries.

| Income category | Median factor increase to national budgets | Median budget increase as % GDP | Median factor increase in conserved land | Median factor increase in conserved ocean | Total budget increase (vs. current spending) \$billions |
| --- | --- | --- | --- | --- | --- |
| HIC | 1.77; 1.83 | 0.091; 0.091 | 1.68 | 1.37 | 12.52; 13.05<br>(17.24) |
| LAMIC | 5.53; 5.35 | 0.103; 0.087 | 2.16 | 8.95 | 9.92; 9.56<br>(3.42) |
| LMIC | 5.41; 4.98 | 0.089; 0.079 | 2.38 | 21.75 | 3.78; 3.62<br>(1.05) |
| UpperMid | 6.27; 6.15 | 0.110; 0.109 | 2.00 | 5.59 | 6.15; 5.94<br>(2.37) |
| LowerMid | 4.05; 3.90 | 0.077; 0.071 | 2.37 | 19.79 | 3.04; 2.94<br>(0.87) |
| Low | 6.46; 5.77 | 0.316; 0.271 | 2.40 | 50.46 | 0.74; 0.68<br>(0.18) |

**Figure 1.**
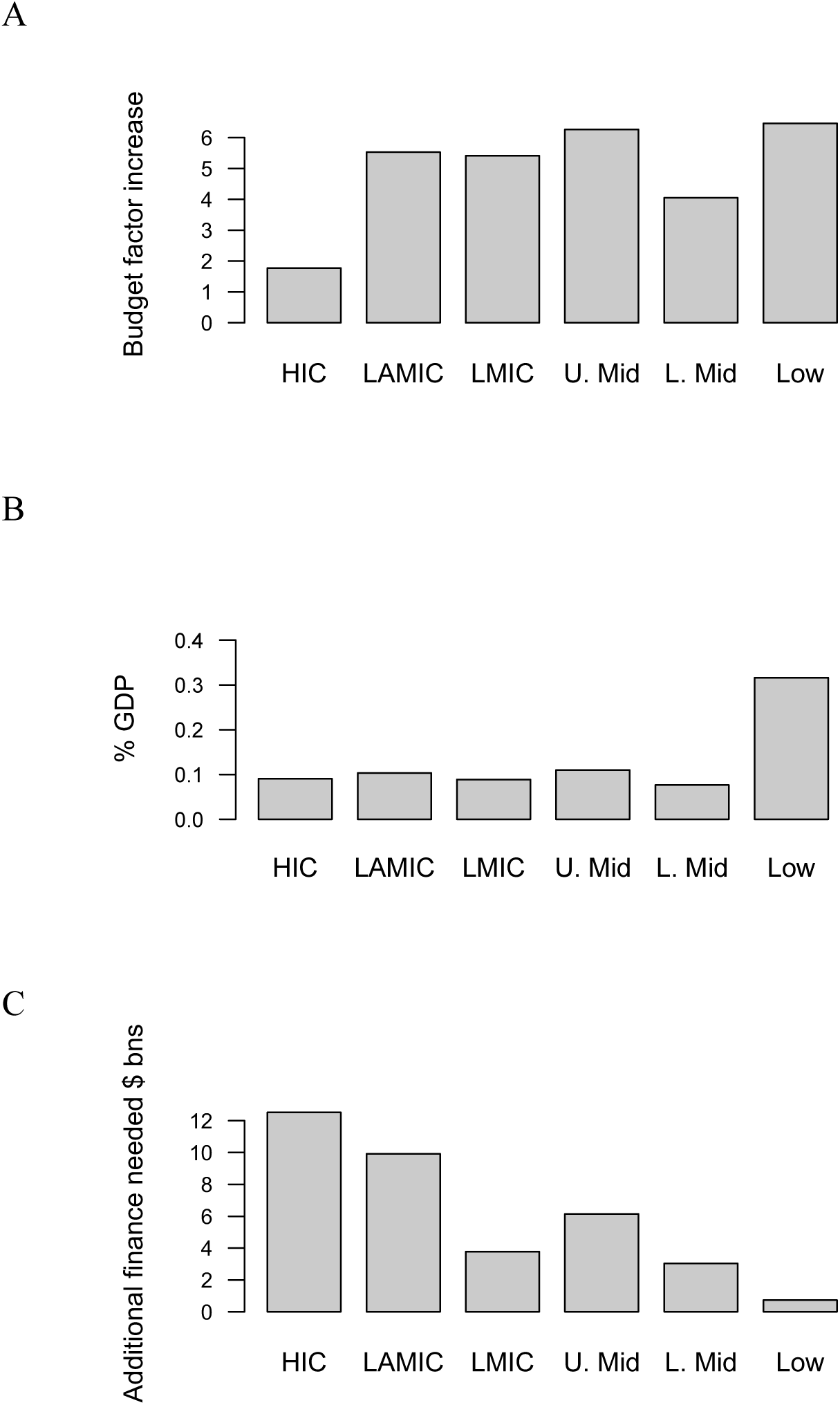
Increase in recurrent annual costs implied by Target 3 (mean of three scenarios). A. Increase expressed as a factor increase (i.e. the Target 3 budget as a multiple of the current budget, median per income group); B. Increase expressed as %GDP (median per income group); C. Raw dollar-value increase in billions of US dollars (total per income group). The baseline scenario for international assistance is shown, see Table 1 for outcomes if aid is doubled. HIC = High-Income Countries, LMIC = Low-to-Middle-Income Countries (Low and Lower-Middle only), LAMIC = LMIC + Upper-Middle-Income Countries (i.e. all non-HIC countries), U. Mid = Upper-Middle Income Countries, L. Mid = Lower-Middle Income Countries.

If current aid to PCAs were doubled, the impact on the distribution of Target 3 costs would be very small (Table 1). For example, a doubling of aid would reduce the budget factor increase in LMICs from 5.41 to 4.98. Doubling aid only makes a small difference because current aid accounts for a relatively small proportion of lower-income PCA expenditures: 5.4% in upper-middle income countries, 7.3% in lower-middle income countries, and 25.9% in low income countries (noting that these values may be overestimates, see Methods).

Under our scenarios for Target 3, lower-income countries would also have to commit more new territory to conservation than HICs (a higher “territorial cost”). On average (mean across the scenarios), HICs committed 1.68 times more new land to conservation, compared with 2.38 times in LMICs and 2.16 times in LAMICs (Figure 2, Table 1). On the sea, the difference was more marked: HICs would need to commit 1.37 times more of their Exclusive Economic Zone (EEZ), LMICs 21.75 times more, and low income countries 50.5 times more (Figure 2, Table 1). These very large increases reflect the current paucity of marine protected areas in lower income countries.

**Figure 2.**
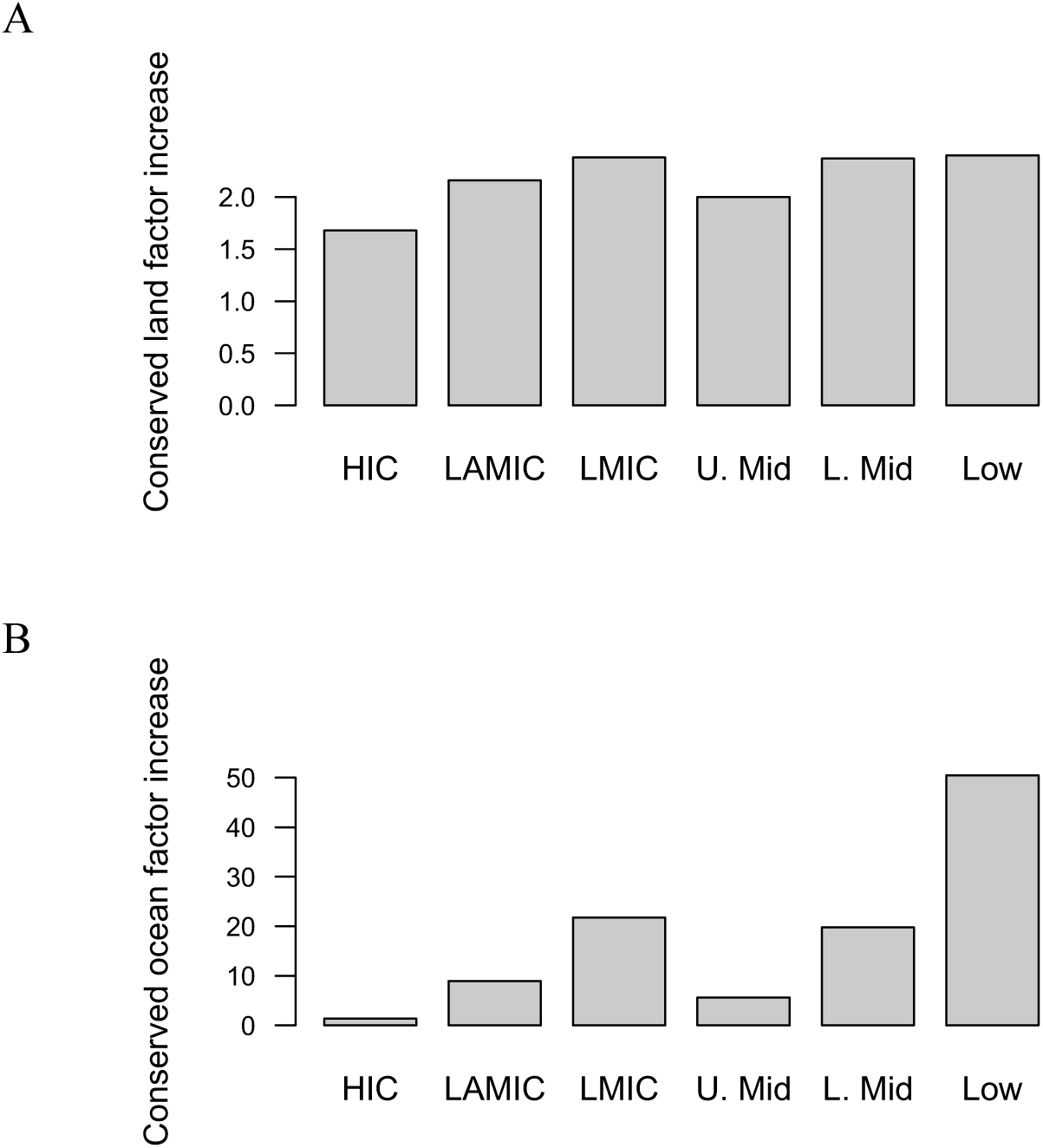
Factor increase in the amount of national territory needed to achieve Target 3 (“territorial cost”). A. On land; B. On the ocean. See Figure 1 for acronyms. Values show the mean across the three scenarios.

## Implications for cost sharing and equity

Our analysis shows that Target 3 would imply a much larger increase in costs for lower-income countries than for high-income countries, both in terms of finance and the amount of national territory committed. Simple, ethical considerations of fairness would suggest that contributions and cost shares should at least be equal and in reality, that HICs might contribute more to global goals, as befits their greater economic capacity. If the disproportionately large burden on lower-income countries is not mitigated, eventual commitments at COP15 could become diluted or underfunded, in ways that threaten the achievement of the underlying biodiversity and climate goals. The disproportionately large burden on lower-income countries seems especially stark when one considers that high-income countries receive a disproportionately large amount of the benefits of biodiversity conservation^12,13^.

We are not seeking to imply that the entire extra burden should automatically fall on wealthier countries. In the long run, developing economies may achieve the (modest) capacity to fund their own PCA systems, as an appropriate investment in preserving the economic and social value of their national natural capital. In terms of simple %GDP, this long-term goal is not infeasible and the values per income group are often similar (Table 1). However, %GDP, for all its popularity as a public-spending metric, is not a good measure of cost-sharing equity across disparate income groups. Governments differ in the proportion of GDP that is available for national public expenditure, with debt burdens particularly reducing this proportion in lower-income countries, and so similar %GDP between income groups can conceal very different realities for finance ministries.

The factor increase is likely to give a better indication of cost-sharing equity, since current spending (the baseline of the factor increase) broadly reflects available public budgets. In the short term (and for COP15), the factor increases suggest that the additional financial burden on lower income countries seems both excessively large and unfairly distributed, particularly for countries that are already using up to 40% of government resources for foreign debt repayments^14^. Some increase in international assistance, innovative finance (including climate finance directed to nature-based solutions^15–17^), or other international finance flow is therefore suggested. Concerningly, we found that international assistance to PCAs currently represents a very small proportion of the budget need, and so even a 100% increase in assistance would have a trivial impact on equitability (a small amount doubled is still a small amount). Indeed, if the LMIC/LAMIC area under conservation will more than double under Target 3, then merely doubling assistance would seem like a step backwards on equitable burden-sharing. Much higher international funding would likely be needed, potentially implying a structural change to the way donors typically support projects rather than recurrent annual expenditure in protected areas.

Upper-Middle Income countries (UMICs) present a serious and separate cause for concern, with further implications for structural change. Several UMICs combine extremely high biodiversity or forest cover with rapidly increasing pressure from their developing economies (e.g. Malaysia, Mexico and South Africa), making it urgent that they expand their PCA systems. However, their higher cost bases also make PCAs more expensive, as illustrated by the five-fold budget increase that they may face for Target 3. And yet, UMICs receive little international aid in current aid allocations. We acknowledge that some UMICs could contribute more to PCAs and biodiversity and indeed, some have been increasing their budgets^18,19^. But even so, such disproportionately large cost increases seem unreachable in the short term. A revision of the patterns of international funding may therefore be called for, perhaps revisiting the way aid allocations taper sharply for UMICs; analysing how far domestic UMIC contributions could feasibly increase; and exploring other funding mechanisms such as carbon payments, green/blue loan strategies and debt swaps^20^.

We caution that our calculations consider only the distribution of recurrent annual management costs and not the distribution of the one-off PCA creation costs. Although this omission is necessary because creation costs vary in unpredictable ways across income groups (see Methods), the annualized cost of purchasing new land for conservation could increase the annual expenditure needs by at least a factor of two^21^, further stressing budgets in lower-income countries. We also caution that the budget needs calculated here are based on a “basic” funding-adequacy scenario. For PCAs to achieve their goals fully (the “optimal” level of funding), budgets would need to be increased further, although there is less data on this. Where countries did report an optimal budget need for PCA management, it was on average 180% of the “basic” need in LAMICs^3^, with the percentage increase for HICs unknown but likely to be less^22^. If Target 3 is funded optimally rather than at a “basic minimum”, the inequalities would therefore be even larger than we found here. If opportunity costs in lower-income countries also increase more than in high income countries, as the territorial cost analysis suggests, then even greater inequity of cost sharing would occur.

We have not attempted to define exactly what an equitable cost distribution would look like, because equity is not a scientific question but a legal and moral one (and in negotiations, ultimately a political one). Legally, states enjoy differential treatment under international law due to their different economic, social and historical situations^5^. Shared actions on global goals should follow a principle of equity^5^ and for biodiversity specifically, Principles 6 and 7 of the 1992 Rio Declaration state that countries have differing circumstances and bear differing responsibility for global environmental problems (see e.g. Sumaila et al.^23^), and so will need to make differing contributions to the solution^24^. A fully equitable arrangement for cost-sharing (burden sharing) would also logically reflect the distribution of benefits. For example, HICs derive many benefits from protecting biodiversity in lower income countries, and a disproportionately high share of benefits overall: examples would be the benefit of climate stability if tropical forests are better conserved, or the value that inhabitants of HICs derive from experiencing species and landscapes in other countries^12^.

We acknowledge that many countries and commentators use the phrase “common but differentiated responsibility” (CBDR) to refer to differential treatment^24^ and CBDR has indeed been extended to refer (among other things) to the distribution of financial commitments for REDD+ ^25^. However, CBDR is not explicitly included in the CBD and remains controversial^26^, as it was for climate talks until it was finally incorporated into the UNFCCC process in 2012^27^. It is beyond the scope of this article to describe the CBDR debate. However, we comment that it is not necessary for parties to formalize CBDR in order to accept the simple, general principle that countries differ in their ability to fund global environmental goals, and that some countries may legitimately need to contribute more than others, if global agreements are to be reached.

We do not address whether the 30% goal is sufficient to achieve the underlying 2050 mission of halting biodiversity decline. If more conservation area is needed, then the cost-sharing inequities could be worse than those shown in our results. We also suggest that other parts of the GBF, and indeed of the Sustainable Development Goals and UNFCCC, should have similar analyses of their cost distributions, as multiple cases of inequitable burden-sharing, across different individual targets, could further threaten the achievement of broader international goals.

Finally, it is worth noting that the absolute budget increase needed in LMICs and LAMICs represents less of a potential financial commitment for donor countries than might be imagined. For example, our model suggests that the additional Target 3 recurrent costs for all LAMICs might be about $9.9 billion per year, equivalent to 0.018% of the total GDP of all high-income countries. In the future, all countries may achieve the capacity to fund the conservation of their own natural capital but in the interim, a mixture of both domestic funding and international aid is likely to be needed. Failing to commit these fairly small amounts of equitable nature-protection funding would simply be a false economy, costing much more than the amount briefly saved, as biodiversity loss and climate crises continue to damage global economic output^12,28,29^.

## Methods

We considered the financial costs of meeting draft post-2020 CBD Target 3 in terms of the national budgets needed for recurrent annual management of an expanded PCA system. To calculate a likely range of costs, we took three of the scenarios created for an existing study on a 30% coverage ambition (Waldron et al. 2020^3^). The original study used six scenarios but not all of those scenarios correspond to the language now being used for Target 3 (in particular, the focus on areas of importance for biodiversity^1^). Our scenarios were specifically as follows: Scenario 1, “Biodiversity Focus”, uses an algorithm to select 30% of the land and sea area to be protected based on its biodiversity importance. Scenario 3, “Harsh Political Reality”, does not allow PCAs on any areas that will be needed for the most efficient agricultural or fisheries production, and then conserves the most biodiversity-important 30% that remains after those production areas have been masked out. Scenario 2, “Biodiversity-Production Compromise”, is intermediate between those positions: on land, it prioritizes biodiversity for the first 5% of additional PCA area (expanding on the existing protected area system), then applies the Harsh Political Reality principle for the remaining land needed to reach the 30% target. On the oceans, it models permission for sustainable fishing on half of the protected marine 30% and no-exploitation on the other half. (See Waldron et al. 2020^3^ for a full description of the cost algorithms and scenario creation methods.)

We also used updated versions of the costing algorithms developed in the same report^3^. Full details can be found in the original source but to summarise: for terrestrial protected and conserved areas, we developed a statistical model based on the reported funding needs of national protected area systems across lower-income countries, and supplemented this with a broad estimate of the funding shortfall for protected areas in High Income Countries based on a recent study by Besancon et al.^22^. Statistical models were able to predict national costs per hectare with a high degree of accuracy (R^2^ = 0.87), based on the size of the protected area system, the mean human footprint^30^ and net agricultural rent^31^ in the areas surrounding the protected areas (expressed relative to the mean human footprint and mean net agricultural rent in the entire country), government effectiveness (i.e. quality of governance), and the level of site-based revenue per hectare (i.e. many protected areas operate as businesses and the higher the revenue, the higher the costs associated with capturing that revenue). For marine protected areas (MPAs), a similar dataset was collated, and costs per hectare were predicted (with R^2^ = 0.90) using a statistical model containing, as predictor variables, the size of the national marine protected area estate, the mean GDP per capita in the zones surrounding the protected areas (adjusted for distance offshore), and the ratio between the number of international tourist arrivals and the population of the country (as a proxy for potential site-based revenue or similar tourism effects). Our updated marine models also accounted for the way expanded MPA systems are likely to include considerably more offshore areas, which are likely to require remote patrolling assisted by satellite tracking and remote electronic monitoring of fishing vessels, where these offshore management methods were assumed to have a different cost structure from the more near-shore management methods that the statistical model was parameterized upon.

To project likely recurrent annual costs for Target 3, these models were then applied to new sets of data that reflected the change in variables expected from the three scenarios of how Target 3 might be implemented (see main text and Waldron et al. 2020 for descriptions of the scenarios themselves). All costs were analysed at purchasing power parity but were back-converted to US dollars at 2015 constant values for reporting. To disaggregate these results by income group, we took the 2021 World Bank income group classification. We note that the models are not able to calculate cost estimates for the overseas territories of HICs directly (due to the way global economic statistics focus on sovereign countries); overseas territory costs are therefore omitted and the HIC costs for countries with such territories will be underestimated as a result, although the size of the territory-specific underestimate is a very small fraction of the total “homeland” budget for those countries, and therefore has a trivial impact on our overall results. Calculating budget increases requires an estimate of current spending, which we also took (and updated) from Waldron et al^3^.

We calculated annual PCA management budget increases for Target 3 as factor increases (multiples of current values) and as %GDP; in both cases, we took the median of the national outcomes for those values in each income group. Increases in territorial costs (the land or sea committed to conservation in PCAs) were calculated at the aggregate level of income groups, rather than the median % per country, due to several countries having either no terrestrial or no marine protected areas formally registered at present in the World Database on Protected Areas^32^. We did not attempt to disaggregate the distribution of PCA creation costs across income groups, because we cannot define how those groups might differ in the amount of PCA land that needs to be purchased (versus how much needs no purchase e.g. because it is state-owned land or an OECM).

To account for the fact that some PCA expenditure in lower-income countries will be met by aid from HICs, we also used our empirical cost data^3^ to estimate the median proportion of expenditure on national PCA systems that is currently met by aid in each income group (median data year = 2015). The aid portion of expenditure was subtracted from the funding commitments for the lower-income groups and added to the commitments for HICs. We also used the empirical aid values to estimate the cost distributions if aid is doubled as part of the implementation of Target 3. We caution that the majority of the data on aid proportions comes from countries that have been recipients of donor assistance, particularly from the Global Environment Facility, and so the aid proportions we use may be overestimates (because countries that did not report aid assistance for their PCA budgets are likely to have lower proportions of assistance and potentially, zero assistance).

We emphasize that, for protected areas, spending data and budget need data are not always consistently estimated and are difficult to collate with a high degree of accuracy, and modelled results are also subject to error. The values we report here for budgets and their distributions should not be interpreted as exact, although they are likely to capture the broader pattern of what the costs and distributions are likely to be. We also caution that the Target 3 costs reported here are not comparable to the costs reported in Waldron et al. 2020 for a 30% goal because the costs in this paper average across a different set of scenarios; omit annualized creation costs; and are based on a set of updated statistical models and approaches.

## Acknowledgments

The work received funding support from National Geographic and the Resources Legacy Fund. D.C. Miller commented on the manuscript.

## Notes

### Competing Interest Statement

The authors have declared no competing interest.

